# Faces and bodies are increasingly integrated along the visual hierarchy in humans and deep neural networks

**DOI:** 10.64898/2026.02.16.706115

**Authors:** Leonard E. van Dyck, Katharina Dobs

## Abstract

Human visual cortex contains regions specialized for faces and bodies, yet we perceive people as a whole. Why does the brain appear to segregate faces and bodies, and how are they integrated to support person perception? Here, we test whether deep neural network models optimized for visual recognition develop segregated or integrated face and body processing, and how this compares to fMRI activity in visual cortex during natural image viewing. We find that models contain face- and body-selective units but also mixed-selective units that are tuned to both faces and bodies. While face- and body-selective units explain unique variance in their corresponding cortical regions, mixed-selective units best explain activity across regions, and shared variance increases from posterior to anterior cortex. Together, our findings suggest that face and body processing reflects a balance of segregation and integration along the visual hierarchy in humans and models, supporting specialized yet flexible person perception.

## Introduction

The perception of other people is a core function of the human visual system. We recognize individuals, read emotions, and infer intentions from faces and bodies. To support these diverse abilities, visual cortex must extract distinct facial and bodily features and combine them into coherent percepts of whole persons. Neuroimaging studies have revealed face- and body-selective regions in high-level visual cortex that respond preferentially to faces or bodies over other categories (Freiwald et al., 2016; Kanwisher, 2010; Peelen & Downing, 2007; Pinsk et al., 2009). Yet these regions are commonly adjacent and often overlap, blurring their functional boundaries (Kim et al., 2014; Kliger & Yovel, 2020, 2024; Peelen & Downing, 2005; Schwarzlose et al., 2005; Weiner & Grill-Spector, 2010, 2013). This raises a fundamental question: To what extent are face and body processing segregated or integrated in visual cortex?

Current theories span a broad spectrum (Taubert et al., 2022). A fully segregated account holds that faces and bodies are processed in distinct functional pathways (Moeller et al., 2008; Pitcher et al., 2009; Premereur et al., 2016; Schwarzlose et al., 2005; Tsao et al., 2006), and that apparent overlap primarily reflects methodological limitations, such as fMRI’s spatial resolution. In contrast, a fully integrated account proposes that faces and bodies share a continuous representational space (Haxby et al., 2011, 2020; Huth et al., 2012; Tarhan & Konkle, 2020), and that apparent separation mainly arises from stimulus or analysis choices, such as cropped images or univariate contrasts. Intermediate, partly integrated accounts posit that processing is segregated in posterior regions but integrated in anterior regions (Bernstein et al., 2014; Harry et al., 2016; Hu et al., 2020; Liu et al., 2010; Taylor et al., 2007). Moreover, a stronger version of this account suggests that integration emerges early on and increases progressively along the visual hierarchy (Taubert et al., 2022).

Despite the fundamental insights from neuroimaging studies, whether and how visual cortex segregates or integrates faces and bodies remains unclear (Hu et al., 2020; Taubert et al., 2022). Task-optimized deep neural networks (DNNs) offer a computational perspective on this question, providing a systematic framework for testing why certain principles of cortical organization emerge (Cichy & Kaiser, 2019; Doerig et al., 2023; Kanwisher et al., 2023; Kietzmann, McClure, et al., 2019; Richards et al., 2019). Recent work has shown that DNNs optimized for visual recognition develop category-selective units that align with category-selective cortical regions, suggesting that selectivity is a hallmark of functional organization in visual systems (Blauch et al., 2022; Janini & Cichy, 2026; Margalit et al., 2024; Prince et al., 2024; Ratan Murty et al., 2021). Task optimization in DNNs segregates information for unrelated tasks such as face and object processing (Dobs et al., 2022) but integrates information for related tasks (Yang et al., 2019). Faces and bodies are visually distinct yet inseparable as parts of the same person, creating pressure for both segregation and integration. How do visual cortex and DNNs balance the competing demands of face and body processing, and do both systems arrive at the same solution?

In this study, we used DNNs to test whether optimization for visual recognition produces segregated or integrated face and body processing. We first applied an fMRI functional localizer (Stigliani et al., 2015) to identify DNN units that respond preferentially to faces, bodies, or both. We then fit encoding models using a large-scale fMRI dataset of natural image viewing (Allen et al., 2022) to examine how each unit type predicts activity in face- and body-selective cortical regions. Each theoretical account makes distinct predictions: (i) A fully segregated account predicts distinct face- and body-selective units that map exclusively to their corresponding regions. (ii) A fully integrated account predicts broadly distributed mixed-selective units that explain different regions similarly well. (iii) A partly integrated account predicts selective units in early layers explaining primarily unique variance in posterior regions, with mixed-selective units in late layers explaining primarily shared variance in anterior regions.

To preview our findings, we identified face-, body-, and mixed-selective units across a diverse set of DNNs. Contrary to a fully segregated account, mixed-selective units best predicted fMRI activity in face- and body-selective regions. Yet contrary to a fully integrated account, face-and body-selective units explained unique variance in their corresponding regions. Consistent with a partly integrated account, shared variance increased gradually from posterior to anterior cortex, reflecting increasing integration along the visual hierarchy. In silico analyses further confirmed that mixed-selective units are functionally relevant for multiple person perception tasks. Our findings reveal that face and body processing in both humans and computational models reflects a principled balance of segregation and integration along the visual hierarchy, supporting specialized yet flexible person perception.

## Results

### Task optimization leads to pure and mixed selectivity for faces and bodies in DNNs

DNNs are widely used as computational models of visual processing and can be systematically varied in architecture, training dataset, and learning objective (Cichy & Kaiser, 2019; Doerig et al., 2023; Kanwisher et al., 2023; Kietzmann, McClure, et al., 2019; Richards et al., 2019). When optimized for visual recognition, do such models develop segregated or integrated processing of faces and bodies? To find out, we analyzed eight DNNs varying in all three dimensions. These included six supervised models spanning three architectures of increasing layer depth (AlexNet, VGG16, and ResNet50) and two training datasets with different category taxonomies (Ecoset and ImageNet), plus two self-supervised models trained on the same datasets using the Barlow Twins objective (Zbontar et al., 2021), which minimizes redundancy between embeddings of differently augmented views of the same image.

We tested whether models develop distinct face- and body-selective units, mixed-selective units, or both. Under a strictly segregated account, face and body signals should be encoded separately by corresponding selective units, whereas under a fully integrated account, they should be encoded jointly by mixed-selective units. To measure selectivity at the finest scale and avoid apparent mixed response profiles caused by spatial pooling, we defined each spatial position within a convolutional layer as a separate unit and analyzed all convolutional and fully connected layers, following previous work (Baek et al., 2021; Lu & Wang, 2025; Prince et al., 2024). We classified units into three types based on their responses to grayscale images of faces, bodies, scenes, and objects, taken from a standard fMRI functional localizer (Fig. 1A; Stigliani et al., 2015). We adapted an approach from previous fMRI studies to identify mutually exclusive selective populations (Kliger & Yovel, 2020, 2024). Face-selective units (hereafter simply “face units”) responded more to faces than to other categories, body-selective units (“body units”) responded more to bodies than to other categories, and mixed-selective units (“mixed units”) responded more to both faces and bodies than to other categories, but did not need to respond equally to both (see Materials and methods).

**Fig. 1.**
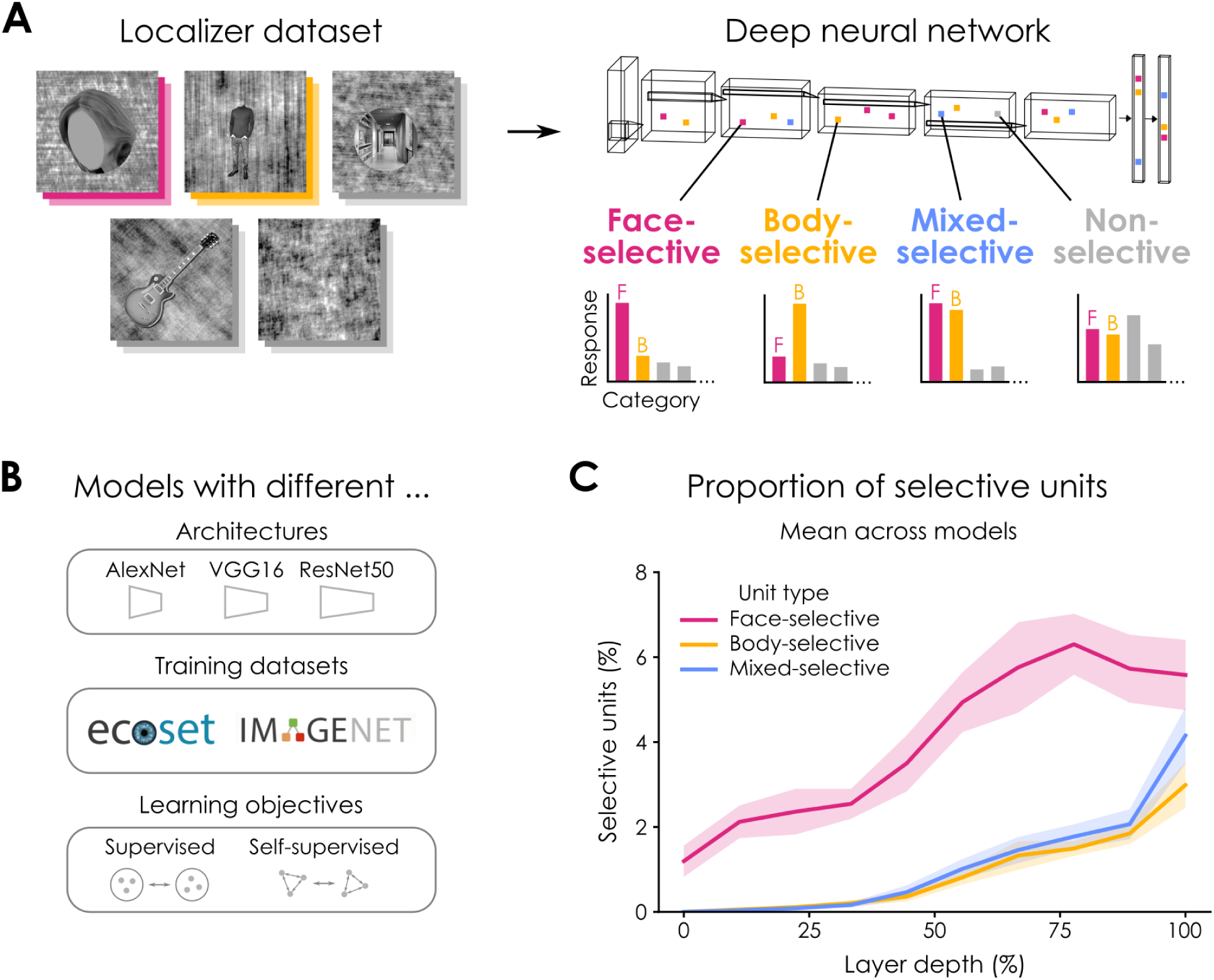
Identifying selective units in DNNs. (**A**) Functional localizer procedure for classifying DNN units as either face-, body-, mixed-, or non-selective. Images containing faces are masked. (**B**) Models (*N* = 8) with different architectures (AlexNet, VGG16, and ResNet50), training datasets (Ecoset and ImageNet), and learning objectives (supervised and self-supervised). (**C**) Proportion of selective units (%) across normalized layer depth (%), averaged across models (mean ± *SEM*).

Across models, we found distinct sets of face and body units alongside a set of mixed units (Fig. 1B). All three unit types became more prevalent across layers (Spearman *r* between layer depth and mean % units across models: face: *r* = 0.89, *p* = 0.001; body: *r* = 1, *p* < 0.001; mixed: *r* = 1, *p* < 0.001), consistent with later layers encoding higher-level category information. Mixed units emerged in intermediate layers, suggesting that faces and bodies begin to integrate based on mid-level features. Face units were most prevalent, while body and mixed units appeared in similar quantities (mean ± *SD* % units in penultimate layer across models: face = 5.58 *±* 2.51, body = 2.99 *±* 1.62, mixed = 4.15 ± 2.01). Given that the criterion for mixed selectivity was more stringent than for body selectivity, their similar prevalence suggests that mixed selectivity is a meaningful functional property rather than an epiphenomenon. Moreover, untrained DNNs also contained units meeting these selectivity criteria (Baek et al., 2021; Prince et al., 2024), but unit counts were uniformly low across layers, and mixed units were nearly absent (Fig. S1).

To determine whether mixed selectivity is specific to faces and bodies or a general property of category representation in models, we examined additional category pairs using the same selectivity criteria. Mixed units also emerged for faces and scenes as well as for bodies and scenes, although at substantially lower rates, particularly in late layers (Fig. S2). This suggests that while mixed selectivity may arise for any pair of related categories, it is especially pronounced for faces and bodies, likely reflecting their frequent co-occurrence in natural images.

These results show that face, body, and mixed units emerge reliably across architectures, training datasets, and learning objectives, and become increasingly prevalent toward later layers. The emergence of mixed units in intermediate layers suggests that face and body signals begin to integrate based on mid-level visual features.

### Selective units encode distinct and generalizable visual features

Having identified different types of selective units using the functional localizer, we tested whether their selectivity profiles generalize to novel images. We evaluated unit responses to faces, bodies, and objects from an independent validation dataset (Fig. 2A), using *d’* to quantify the preference for faces or bodies over objects (see Materials and methods). All unit types showed robust selectivity profiles that generalized to novel images. Face and body units showed a clear double dissociation, with each type responding selectively to its preferred category, while mixed units responded selectively to both. These profiles remained consistent across layers (Fig. S3A). In untrained DNNs, unit types failed to generalize to novel images (Fig. S3B), confirming that robust selectivity depends on learning.

**Fig. 2.**
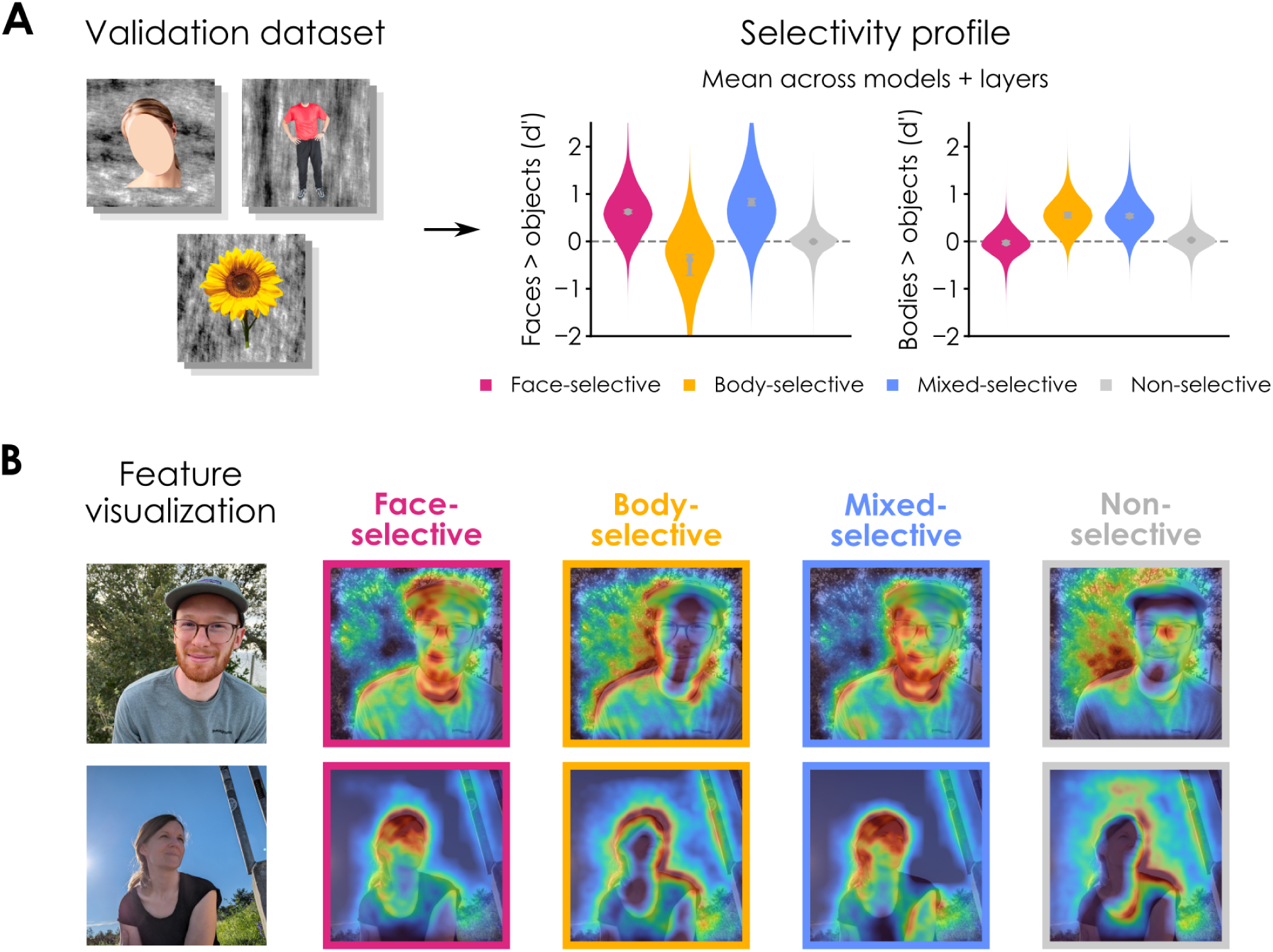
Validating selective units in DNNs. (**A**) Selectivity profiles for each unit type based on an independent validation dataset. Mean face and body selectivity (*d’*) of unit types, averaged across models and layers. Dots and error bars show mean and 95% CIs (hierarchical bootstrapping; 1,000 iterations). Images containing faces are masked. (**B**) Feature visualizations using Guided-GradCAM for each unit type in the last convolutional layer of a representative model (supervised AlexNet trained on Ecoset; conv5). Images containing faces show the authors.

This generalization suggested that units are driven by reliable visual features. To identify the features driving the different selectivities, we applied Guided Grad-CAM (Selvaraju et al., 2017), which combines gradient-based sensitivity with class activation mapping. Feature visualizations for a representative model (supervised AlexNet trained on Ecoset) revealed distinct preferences (Fig. 2B, Fig. S4). Face units responded primarily to facial features and body units to body contours. Notably, body units often also responded to head contours within whole-person images despite being identified using headless body stimuli. Mixed units responded to both facial and bodily features within the same images.

### Mixed units best predict activity in face- and body-selective cortex

Having established face, body, and mixed units in models, we next asked how they map onto face- and body-selective regions of visual cortex. This provided a critical test of segregated versus integrated processing. Under a segregated account, face and body units should best predict activity in their corresponding regions, whereas under an integrated account, the different unit types should show comparable predictive power across regions.

We used the Natural Scenes Dataset (Allen et al., 2022), which contains high-resolution fMRI responses from eight participants viewing thousands of natural images. Functional regions were defined using the same functional localizer applied to models but with standard fMRI contrasts (i.e., target category vs. all other categories, *t* > 2; Fig. 3A). These regions included three face-selective regions—occipital face area (OFA), fusiform face area (FFA), and an anterior temporal face area (aTL-faces)—and three body-selective regions—extrastriate body area (EBA), fusiform body area (FBA), and a medial temporal body area (mTL-bodies). To disentangle pure and mixed selectivity, we excluded voxels selective for both faces and bodies from all regions and defined a dedicated overlap region between FFA and FBA for these mixed-selective voxels, which is commonly observed in fMRI studies (Kim et al., 2014; Kliger & Yovel, 2020, 2024; Peelen & Downing, 2005; Schwarzlose et al., 2005; Weiner & Grill-Spector, 2010, 2013).

**Fig. 3.**
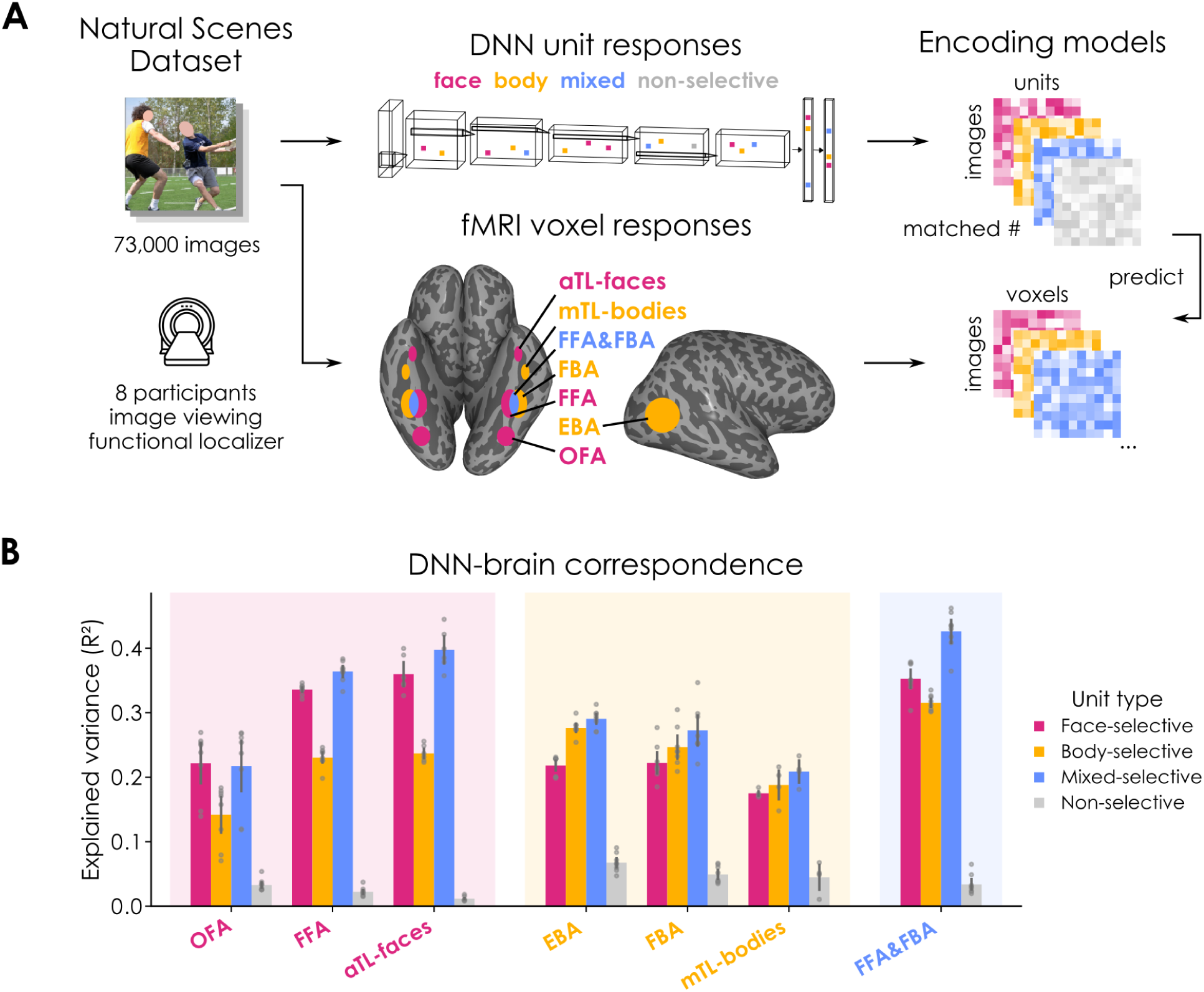
Linking DNN units to cortical regions. (**A**) For a representative model (supervised AlexNet trained on Ecoset), voxel-wise encoding models were fit using non-negative ridge regression for each participant, region, layer, and unit type. Face-selective regions (magenta), body-selective regions (yellow), and commonly observed spatial overlap (blue) are shown on an inflated cortical surface. Images containing faces are masked. (**B**) Bars show mean explained variance (*R^2^*, noise-normalized) in the penultimate layer (fc7, post-ReLU), averaged across voxels and participants. Dots and error bars show participant means and 95% CIs (Cousineau-Morey-corrected).

We fit voxel-wise encoding models with cross-validation to predict voxel responses from unit responses. Ridge regression was used to handle the high dimensionality of the predictors and prevent overfitting. A non-negativity constraint was imposed on the regression weights to aid interpretability, ensuring that each unit type contributed positively to the predicted response. To allow fair comparisons, unit counts were matched across types within each layer by selecting the most selective units of each type based on the minimum count. These subsampled units maintained robust selectivity (mean *d’ ± SD*; face = 1.64 ± 0.23; body = 0.75 ± 0.04; mixed = 0.87 ± 0.22). For each participant, region, and unit type, we computed the mean explained variance (*R^2^*) across voxels. To compare unit types within each region type (i.e., face-selective, body-selective, or overlap), we used pairwise *t*-tests across participants (two-sided, *p* < 0.05, Bonferroni-corrected).

Units in the penultimate layer consistently provided the best predictions, explaining voxel responses well relative to the estimated noise ceiling across all regions (Fig. 3B). These units also outperformed non-selective units throughout (all *p* < 0.001), consistent with encoding higher-level person information. Mixed units outperformed face and body units in face-selective regions (mixed vs. face: *t*(7) = 5.63, *p* = 0.005, *d* = 1.99; mixed vs. body: *t*(7) = 27.96, *p* < 0.001, *d* = 9.89), body-selective regions (mixed vs. face: *t*(7) = 10.44, *p* < 0.001, *d* = 3.69; mixed vs. body: *t*(7) = 4.61, *p* = 0.015, *d* = 1.63), and the overlap region (mixed vs. face: *t*(7) = 13.18, *p* < 0.001, *d* = 4.66; mixed vs. body: *t*(7) = 12.41, *p* < 0.001, *d* = 4.39). However, mixed units did not outperform face and body units in OFA and EBA (OFA: mixed vs. face: *t*(7) = -0.81, *p* = 1, *d* = -0.28; EBA: mixed vs. body: *t*(7) = 2.37, *p* = 0.297, *d* = 0.84), consistent with these regions representing earlier stages of face and body processing (Harry et al., 2016; Hu et al., 2020; Taubert et al., 2022). Notably, face and body units demonstrated a clear double dissociation, as face units outperformed body units in face-selective regions (face vs. body: *t*(7) = 20.74, *p* < 0.001, *d* = 7.33), while body units outperformed face units in body-selective regions (body vs. face: *t*(7) = 5.72, *p* = 0.004, *d* = 2.02). This pattern held across regions and layers, with mixed and preferred-category units consistently outperforming other-category and non-selective units (Fig. S5). The only exception was mTL-bodies, identified in just half of the participants, where predictive performance did not differ significantly between unit types. As a control, selective units performed poorly in early visual cortex (V1), where only non-selective units from intermediate layers showed moderate performance, confirming the specificity of these effects.

### Segregation and integration are balanced along the cortical hierarchy

Having established that mixed units best predict activity across face- and body-selective regions, we next asked whether face and body units explain unique or shared variance within these regions, and whether this balance shifts along the cortical hierarchy. Under a segregated account, face and body units should each explain mostly unique variance in their corresponding regions. Under an integrated account, their contributions should largely overlap.

To test these predictions, we used commonality analysis, which decomposes the explained variance of a joint encoding model into components unique to each predictor set and components shared between them. We fit joint encoding models using face and body units combined, partitioning their contributions into variance unique to face units, unique to body units, and shared between them. Face and body units explained substantial shared variance across all regions, but each also contributed unique variance in their corresponding regions (Fig. 4A, Fig. S6), consistent with partly segregated processing. To test whether mixed units carry additional information beyond face and body units, we added them to the encoding models. Mixed units explained little additional unique variance, consistent with encoding a combination of face and body signals.

**Fig. 4.**
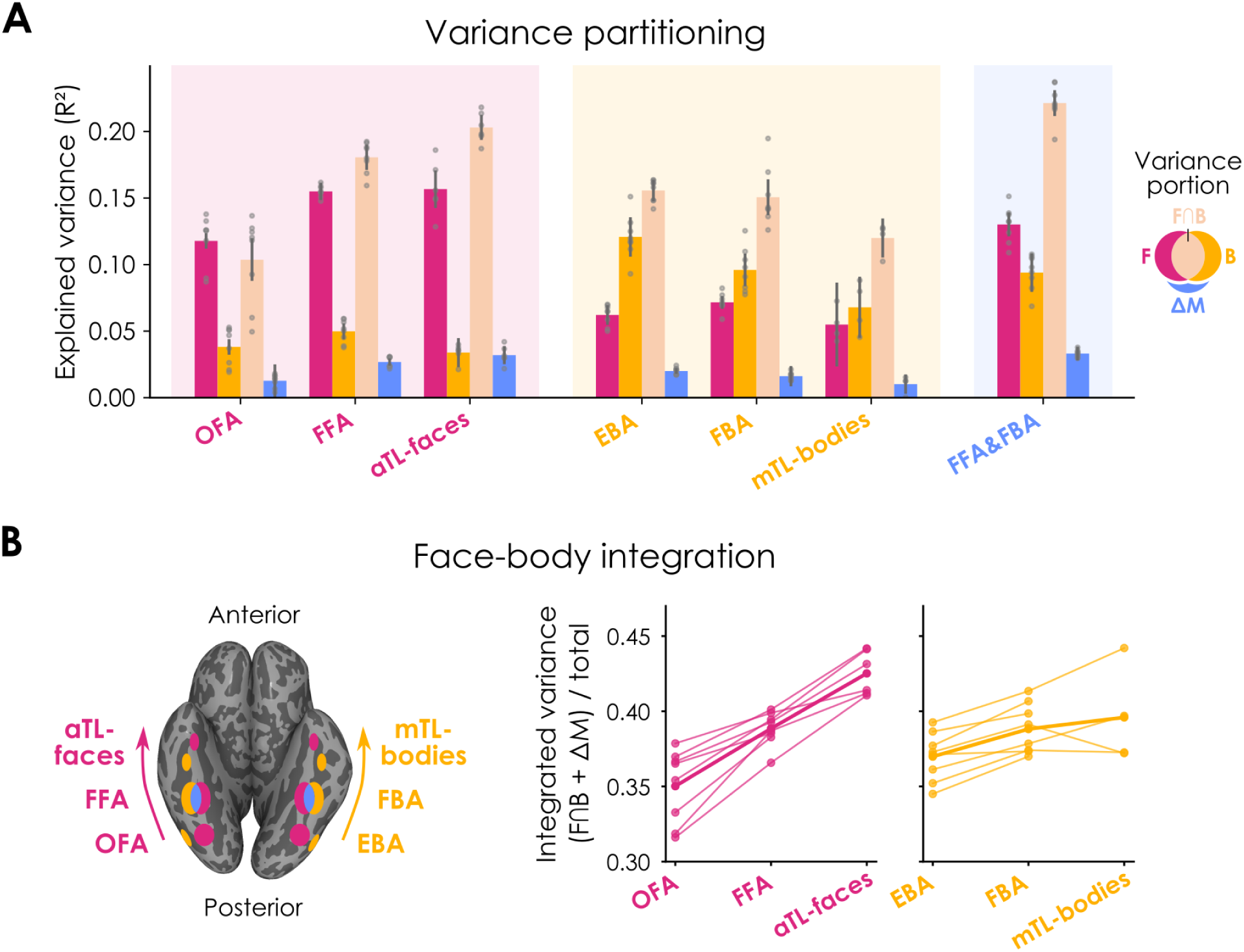
Variance partitioning of DNN unit contributions in cortical regions. (**A**) Commonality analysis of joint encoding models, showing variance explained uniquely by face units (magenta), uniquely by body units (yellow), shared between face and body units (orange), and additionally explained by mixed units (blue). Bars show mean explained variance (*R^2^*, noise-normalized) in the penultimate layer (fc7, post-ReLU), averaged across voxels and participants. Dots and error bars show participant means and 95% CIs (Cousineau-Morey-corrected). (**B**) Face-selective (magenta) and body-selective (yellow) hierarchies are shown on an inflated cortical surface. Integrated variance, defined as the fraction of total explained variance attributable to shared face-body contributions and unique mixed contributions (F∩B + ΔM), across the cortical hierarchy. Thin lines show individual participant means and thick lines show group means.

To examine how this balance shifts along the cortical hierarchy, we quantified integrated variance as the fraction of total explained variance attributable to shared contributions from face and body units plus unique contributions from mixed units. Integrated variance increased systematically from posterior to anterior regions in both face-selective and body-selective cortical hierarchies (Fig. 4B), confirmed by linear mixed-effects models (face: *β* = 0.04, *z* = 11.35, *p* < 0.001; body: *β* = 0.01, *z* = 4.10, *p* < 0.001). These results reveal a gradient from partly segregated processing in posterior regions to increasingly integrated processing in anterior regions.

Together, these results show that mixed units predict activity in face- and body-selective cortex better than face and body units, particularly in anterior regions. Moreover, they reveal a systematic gradient along the visual hierarchy, as posterior regions maintain partly distinct face and body representations whereas anterior regions converge toward increasingly integrated whole-person representations.

### Mixed selectivity supports flexible person perception

Why do models develop units that respond to both faces and bodies? Although such mixed units explained little unique variance in visual cortex, examining their functional role may reveal the principles driving face-body integration in models and possibly in the brain.

To test this question, we trained linear classifiers on unit responses from the penultimate layer for three person perception tasks. For face recognition, we used VGGFace2 (Cao et al., 2018), containing face-cropped images of 500 identities. For person recognition, we used Celeb-reID (Huang et al., 2020), containing person-cropped images of 632 identities across varied clothing, poses, and backgrounds. For action recognition, we used Stanford 40 Actions (Yao et al., 2011), containing person-cropped images of 40 actions (e.g., running, climbing, making a phone call) performed by different individuals. All classifiers achieved robust performance on held-out test sets (face recognition top-5 = 55.17%, person recognition top-5 = 18.19%, action recognition top-1 = 40.94%), confirming that models optimized for visual recognition contain linearly decodable information for diverse person perception tasks. To quantify the functional contribution of each unit type, we performed lesioning analyses (Fig. 5A). For each unit type, we lesioned matched numbers of the most selective units by setting their activations to zero throughout all layers of the fixed DNN backbone and measuring the resulting accuracy drop. We repeated this across ten stratified splits and estimated uncertainty using hierarchical bootstrapping (see Materials and methods).

**Fig. 5.**
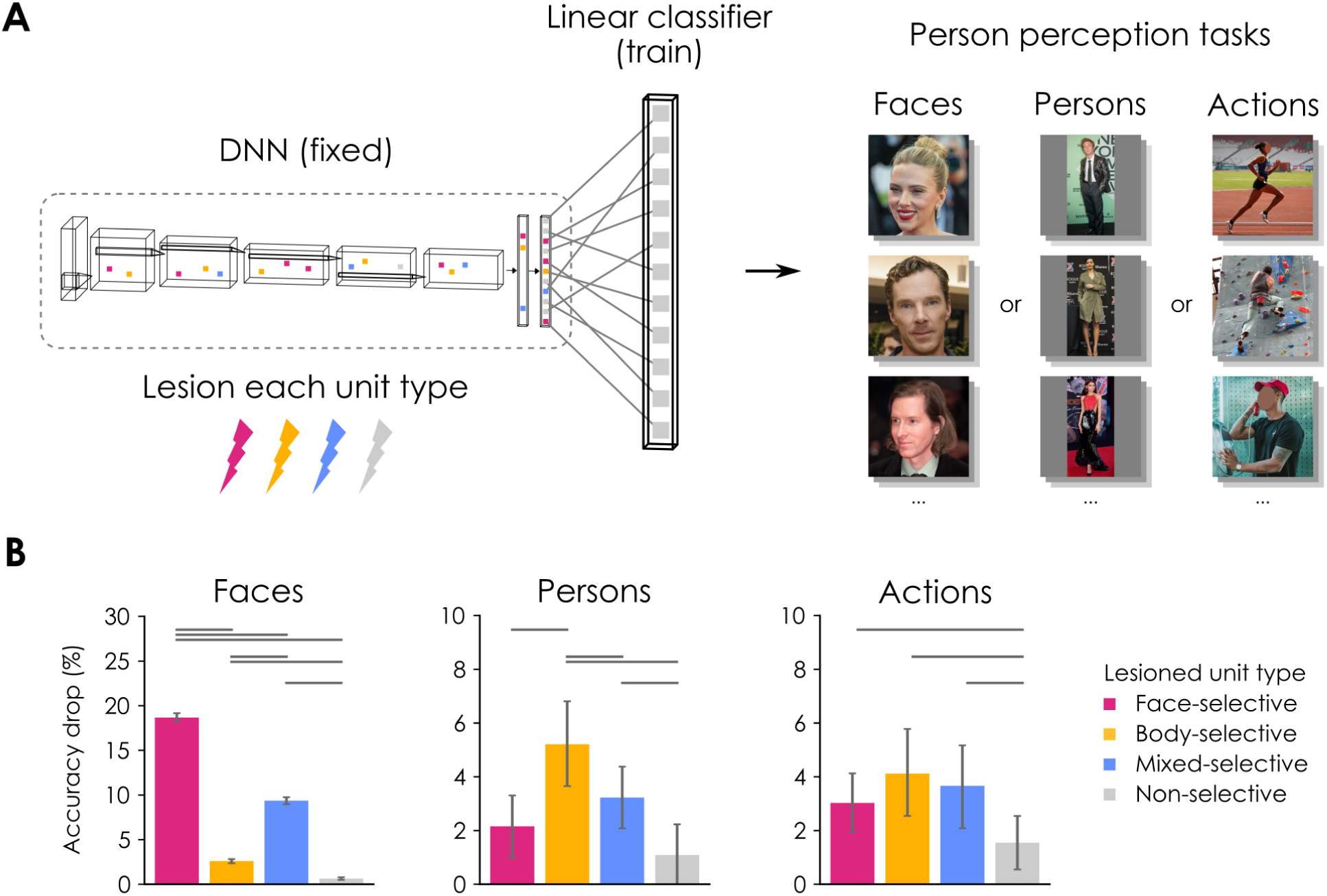
Impact of lesioning unit types on person perception tasks. (**A**) For a representative model (supervised AlexNet trained on Ecoset), linear classifiers were trained on unit responses from the penultimate layer (fc7, post-ReLU) to solve three person perception tasks: face recognition (VGGFace2), person recognition (Celeb-reID), and action recognition (Stanford 40 Actions). After training, a matched number of units of each type were lesioned, and the resulting accuracy drop was measured. For copyright reasons, example images were replaced (sources: Wikimedia Commons and Pixabay.com). Images containing faces of private individuals are masked. (**B**) Normalized accuracy drop (%) for each task after lesioning each unit type. Error bars show 95% CIs (hierarchical bootstrapping; 10,000 iterations). Horizontal bars indicate significant pairwise comparisons (*p* < 0.05, FDR-corrected).

Lesioning selective units impaired performance across all tasks (all *p* < 0.001; Fig. 5B), but the impact varied systematically with task demands. For face recognition, lesioning face units caused the largest deficit (normalized accuracy drop = 18.67%), exceeding mixed units (9.37%; *ΔM* = 9.29, *p* < 0.001, *d* = 12.6) and body units (2.61%; *ΔM* = 16.06, *p* < 0.001, *d* = 18.94), with mixed units also surpassing body units (*ΔM* = 6.78, *p* < 0.001, *d* = 13.53). This demonstrates that face units are specialized for face identity information, while mixed units also encode substantial identity information despite their broader tuning. For person recognition, this pattern reversed. Lesioning body units caused the largest deficit (5.21%), exceeding both mixed units (3.23%; *ΔM* = 1.98, *p* = 0.009, *d* = 1.07) and face units (2.16%; *ΔM* = 3.07, *p* < 0.001, *d* = 1.20), which did not differ significantly from each other (*ΔM* = 1.06, *p* = 0.122, *d* = 0.74). This indicates that body units are specialized for whole-person identity information, while mixed units provide secondary contributions. For action recognition, lesioning face, body, and mixed units produced comparable deficits (face = 3.03%; body = 4.12%; mixed = 3.66%; all pairwise *p* > 0.15), yet all exceeded non-selective units (all pairwise *p* < 0.01). This suggests that action recognition relies on information distributed across different unit types, rather than on a single specialized population.

These results show that mixed units carry multifaceted information that can be flexibly read out across person perception tasks. Face recognition relies primarily on face units, person recognition on body units, and action recognition on multiple unit types, consistent with more integrative tasks drawing on distributed person-related information.

## Discussion

The extent to which faces and bodies are segregated or integrated in human visual cortex remains actively debated, with neuroimaging studies supporting diverging accounts (Hu et al., 2020; Taubert et al., 2022). By combining DNN modeling with fMRI analyses, we show that face and body processing reflects a mixture of segregation and integration along the visual hierarchy in both humans and computational models.

Across a diverse set of DNNs, we consistently found distinct face and body units alongside a considerable population of mixed units. Face units were most prevalent, consistent with a previously reported tendency toward face selectivity in models (Baek et al., 2021; Prince et al., 2024; van Dyck & Gruber, 2023). This imbalance likely reflects a combination of factors, including greater sensitivity of the functional localizer to fine-grained facial features, biases in training datasets toward close-up portraits, and the privileged status of faces as a stimulus category (Farah et al., 1998; Kanwisher, 2000). Crucially, mixed units emerged in intermediate layers, indicating that faces and bodies begin to integrate in mid-level processing, consistent with evidence that category selectivity in visual cortex is closely tied to mid-level visual features (Jagadeesh & Gardner, 2022; Long et al., 2018; Schluesener et al., 2025).

In visual cortex, we observed a strikingly similar organization, supporting functional alignment between category-selective DNN units and cortical regions (Blauch et al., 2022; Janini & Cichy, 2026; Margalit et al., 2024; Prince et al., 2024; Ratan Murty et al., 2021). Contrary to our expectations, face- and body-selective regions were not best predicted by their corresponding unit types. Instead, mixed units were the strongest predictors across regions, suggesting that these regions encode richer person information than their categorical labels imply. This finding aligns with multidimensional accounts of the functional organization in higher-level visual cortex, which propose that category-selective regions are sparsely tuned to multiple representational dimensions (Contier et al., 2024; Ritchie et al., 2026; van Dyck et al., 2025). Variance partitioning revealed that face and body units each contributed unique variance primarily to their corresponding regions, yet most explained variance was shared across unit types, supporting partly distinct but substantially overlapping face and body representations (Harry et al., 2016; Kim et al., 2014; Schwarzlose et al., 2005). Strikingly, this balance shifted systematically from posterior to anterior regions, extending a previously observed segregation-to-integration gradient to natural images of whole persons (Fisher & Freiwald, 2015; Harry et al., 2016; Hu et al., 2020; Taubert et al., 2022; Zafirova et al., 2022).

What might be the functional advantage of a partly integrated face-body organization? We propose it reflects a balance between specialization and flexibility. Pure selectivity may support fine-grained, category-specific computations, such as recognizing a face identity or reading body pose, while mixed selectivity may support broader computations that extend across categories, such as person recognition or action understanding. Consistent with this view, mixed selectivity has been proposed to yield representations that can be flexibly read out across multiple tasks (Rigotti et al., 2013; Tye et al., 2024). This organization likely emerges from the combined pressures of task optimization for visual recognition (Dobs et al., 2022; G. R. Yang et al., 2019) and the co-occurrence statistics of natural images (Arcaro et al., 2020; Bonner & Epstein, 2021; Kaiser et al., 2019). Because faces and bodies frequently appear together, optimization may favor partly integrated codes that reuse features across categories while preserving sufficient category-specific features to support discrimination. Such an organization could be implemented efficiently through normalization mechanisms that stabilize responses to multi-category inputs (Carandini & Heeger, 2012; Kliger & Yovel, 2020, 2024), eliminating the need for a dedicated neural population for whole persons (Afraz, 2015; Fisher & Freiwald, 2015).

Whether visual cortex represents whole persons differently from the sum of their parts remains debated (Hu et al., 2020), with support for both part-based (Kaiser et al., 2014; Kliger & Yovel, 2020; Zafirova et al., 2022, 2024) and holistic processing (Arcaro et al., 2020; Bernstein et al., 2014; Brandman & Yovel, 2016; Fisher & Freiwald, 2015; Schmalzl et al., 2012). Our findings do not resolve this directly, as interactions between receptive field sizes, nonlinearities, and normalization make it difficult to measure holistic processing in models. Nevertheless, the systematic gradient from segregated to integrated processing in visual cortex and models provides a computational framework within which holistic processing could naturally emerge.

Our study has several limitations worth noting. The DNNs examined here were strictly feedforward, lacking the recurrent and feedback dynamics that shape cortical processing (Kietzmann, Spoerer, et al., 2019; Xie et al., 2025). While mixed selectivity emerged without such dynamics, incorporating them may improve functional alignment. The encoding models remain constrained by fMRI’s spatial resolution, which may obscure finer-scale functional organization if purely selective neurons are interleaved within a voxel (Premereur et al., 2016). However, recent single-unit recordings in macaque visual cortex have reported mixed-selective neurons that integrate face and body information in a configuration-specific manner (Zafirova et al., 2024), closely mirroring our findings and suggesting that mixed selectivity is not merely a spatial artifact.

Our findings point to several directions for future research. Models trained with multiple objectives could reveal how task demands shape the balance between pure and mixed selectivity (Dobs et al., 2022; Driscoll et al., 2024; G. R. Yang et al., 2019). The behavioral relevance of mixed selectivity in humans could be tested in tasks that benefit from integrating face and body cues, such as recognition under challenging conditions or judgments of congruence (Aviezer et al., 2012; Foster et al., 2021; Kliger & Yovel, 2024; Meeren et al., 2005; Rice et al., 2013). Finally, whether this organizing principle extends to the perception of multiple persons (Abassi & Papeo, 2024), or more broadly to other frequently co-occurring categories, remains an open question.

In sum, our findings suggest that face and body processing in human visual cortex and computational models reflects a principled balance of segregation and integration, one that preserves face- and body-specific representations while enabling their flexible combination into whole-person representations.

## Materials and methods

### Deep neural network models

To test generalizability across models, we evaluated eight DNNs varying in architecture, training dataset, and learning objective. Six were trained with supervised learning: AlexNet (Krizhevsky et al., 2012), VGG16 (Simonyan & Zisserman, 2015), and ResNet50 (He et al., 2016), each trained on either Ecoset (Mehrer et al., 2021), which includes explicit person categories (e.g., man, woman, child), or ImageNet (Deng et al., 2009), which contains many images of people but no dedicated person category (K. Yang et al., 2022). The remaining two were self-supervised AlexNet models trained on the same datasets using the Barlow Twins objective (Zbontar et al., 2021), which learns by minimizing redundancy between embeddings of differently augmented views of the same image. Supervised ImageNet weights used default TorchVision implementations, and supervised Ecoset weights were obtained from OSF (https://osf.io/kzxfg/). The self-supervised ImageNet model was obtained from Prince et al. (2024), and the self-supervised Ecoset model was trained following the training parameters and sparse linear readout described in the original work (Prince et al., 2024; Zbontar et al., 2021). To compare layers across architectures, we expressed layer position as a percentage of total depth, binned into ten equal intervals.

### Identifying selective units

To investigate selectivity in models, we analyzed individual units across all convolutional and fully connected layers preceding the final classification layer. In convolutional layers, we defined each spatial position within each filter as a separate unit (rather than pooling across positions), following prior work (Baek et al., 2021; Lu & Wang, 2025; Prince et al., 2024).

We identified selective units using cropped grayscale images of faces, bodies, scenes, and objects (288 images per category) from a standard fMRI functional localizer (Stigliani et al., 2015). We adapted an fMRI approach for identifying face-, body-, and mixed-selective responses (Kliger & Yovel, 2020, 2024) to classify DNN units into three mutually exclusive types: (i) face-selective units preferred faces over other categories, (ii) body-selective units preferred bodies over other categories, and (iii) mixed-selective units preferred both faces and bodies over other categories. Mixed units were not required to respond equally to faces and bodies, only to prefer both over other categories.

We first tested whether units responded more strongly to each category (faces, bodies, scenes) compared to a baseline category (objects) using *t*-tests (one-sided, *p* < 0.001, FDR-corrected using the Benjamini-Yekutieli procedure; Benjamini & Yekutieli, 2001). We then computed the contrast between faces and bodies to determine response preferences. Face-selective units were defined as responding more to faces than objects AND more to faces than bodies AND not more to bodies or scenes than objects (one-sided, *p* > 0.05, FDR-corrected). Body-selective units followed this logic, responding more to bodies than objects AND more to bodies than faces AND not more to faces or scenes than objects. Mixed-selective units responded more to both faces and bodies than objects AND not more to scenes than objects. Finally, all selective units were required to respond more strongly to their preferred category (or categories) than to phase-scrambled images provided in the localizer set. This additional criterion substantially reduced false positives by ensuring selectivity reflected meaningful feature sensitivity rather than low-level image statistics.

In many subsequent analyses, we controlled for unequal group sizes across unit types by ranking units within each type based on selectivity (*d’*, see below) and retaining the top *N* per layer, with *N* set to the minimum number of selective units available across unit types. Non-selective units were sampled by ascending mixed selectivity to provide a baseline with minimal face-body preference.

### Validating selective units

To assess generalization beyond the functional localizer, we used an independent dataset containing cropped color images of faces (150 images), bodies (75 images of headless bodies, 75 images of isolated limbs), and objects (30 categories, 5 images each). Images were obtained from the Hemera Photo Objects database and Bracci et al. (2015). To minimize low-level differences across categories, each cropped image was overlaid on a phase-scrambled background image from the functional localizer set.

To quantify selectivity as a continuous measure, we computed a *d’* index for each unit *u* and category *c* against other categories *o*:

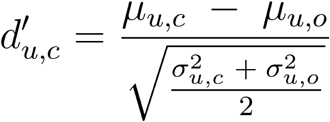

Here *µ_u,c_* and *µ_u,o_* are the mean responses of unit *u* to category *c* and to another category *o*, and *σ^2^_u,c_* and *σ^2^_u,o_* are the corresponding response variances. We computed *d’* using objects as a baseline (face *d’*: faces > objects; body *d’*: bodies > objects). To estimate the mean *d’* of each unit type across models and layers, we used hierarchical bootstrapping (1,000 iterations), as described below.

### Feature visualizations

To visualize features driving each unit type, we used Guided Grad-CAM (Selvaraju et al., 2017). This method combines the coarse spatial localization of Gradient-weighted Class Activation Mapping (Grad-CAM) with the fine detail of Guided Backpropagation (GB) to produce saliency maps indicating both where and what features activate a set of units. All images used in visualizations were obtained with permission or under Creative Commons licenses. For each target layer, we defined a unit-set objective by restricting responses to the selective units of interest and summing their responses into a single scalar score. Spatial localization was obtained with Grad-CAM. For fully connected target layers, the model’s last convolutional layer served as the CAM layer, whereas for convolutional target layers, the preceding convolutional layer was used as the CAM layer. Channel weights were computed as the global average of gradients with respect to CAM activations, and the resulting Grad-CAM maps were upsampled to image resolution and smoothed with a Gaussian blur (*σ* = 5 px). Fine-grained sensitivity was computed with GB saliency maps from input gradients, using ReLU hooks implementing the guided rule (only positive gradients were propagated where forward activations were positive). Robustness was improved by adding zero-mean Gaussian noise to the input (*σ* = 0.05, normalized image space) and averaging saliency maps across 50 repeats. Saliency maps were summed across channels, upsampled to image size, and lightly smoothed (*σ* = 3 px). Final Guided Grad-CAM heatmaps were obtained by element-wise multiplication of the Grad-CAM and GB maps, using weights of 0.7 (Grad-CAM) and 0.3 (GB), which yielded the most interpretable visualizations.

### Natural Scenes Dataset

To investigate selectivity in human visual cortex, we used the Natural Scenes Dataset (NSD; Allen et al., 2022), a large-scale dataset of high-resolution fMRI responses from eight participants. Each participant viewed between 9,000 and 10,000 natural scene images across 30 to 40 scanning sessions, with each image presented up to three times. During scanning, participants maintained central fixation and performed a long-term recognition task that required them to identify previously viewed images. The stimuli were drawn from the Microsoft COCO (Lin et al., 2014) and presented for 3s each at a visual angle of 8.4° × 8.4°, followed by a 1s inter-stimulus interval.

Functional images were acquired at 7T, using a whole-brain, gradient-echo echo-planar imaging sequence, with 1.8mm isotropic voxels and a repetition time of 1.6s. Preprocessing included temporal and spatial interpolation to correct for slice timing and head motion, followed by single-trial response estimation using a general linear model. We used the preprocessed voxel response estimates provided with the dataset, which were optimized through voxel-wise hemodynamic response function (HRF) modeling, data-driven denoising, and ridge regression (“betas_fithrf_GLMdenoise_RR”; Prince et al., 2022). Further details on data acquisition, preprocessing, and noise ceiling estimation are available in the original NSD paper (Allen et al., 2022). To reduce session-specific variability and improve reliability, we *z*-scored voxel responses within each session and averaged the responses across repetitions of each image.

### Functional regions of interest

To identify face- and body-selective cortical regions, the same functional localizer was used in NSD as in our DNN analyses. Participants viewed grayscale images of all 5 categories presented in a miniblock design over 6 runs, with each block containing 8 images of a category. After standard preprocessing, category-specific beta values were estimated with a general linear model, and selective voxels were identified using *t*-tests comparing each category to all others (*t* > 2). Further details are available in the original functional localizer paper (Stigliani et al., 2015). This procedure yielded the following regions: occipital face area (OFA, 8/8 participants), fusiform face area (FFA, 8/8), anterior temporal face area (aTL-faces, 6/8), extrastriate body area (EBA, 8/8), fusiform body area (FBA, 8/8), medial temporal body area (mTL-bodies, 4/8). To isolate pure and mixed selectivity, we excluded voxels selective for both faces and bodies from all regions and defined a dedicated overlap region between FFA and FBA for these mixed-selective voxels (FFA&FBA, 8/8). As a control, we also analyzed primary visual cortex (V1, 8/8) defined using population receptive field mapping. Across all regions, only voxels with a signal-to-noise ratio exceeding 0.15 were included in the analyses.

### Encoding models

To test how well different unit types predict fMRI activity in cortical regions, we used voxel-wise ridge regression with a positivity constraint. The ridge penalty was used to handle multicollinearity and overfitting, while the positivity constraint restricted weights to be nonnegative, enforcing strictly additive contributions (i.e., higher unit activation implies stronger voxel response). To match this constraint, we analyzed unit responses after the ReLU nonlinearity. To ensure fair comparisons, the number of units was matched across types within each layer. To manage computational demands arising from the large number of fits across participants, regions, layers, unit types, and ridge penalty values, we applied non-negative matrix factorization (NMF) to reduce the dimensionality of the matched unit sets. NMF was chosen because it preserves the non-negativity constraint inherent to our approach, ensuring that the reduced components remained interpretable as additive combinations of unit responses. We reduced each unit set to 100 components. In preliminary analyses, encoding performance was highly similar with and without this dimensionality reduction, confirming that NMF retained the predictive information while substantially reducing computational cost.

We trained separate encoding models for each unit type using nested cross-validation. An outer 10-fold split provided unbiased estimates of explained variance (*R^2^*), allowing us to evaluate how well the model predicts new images. Models were trained on 90% of the data and tested on the remaining 10%, repeated across folds. Within each outer training split, an inner 5-fold split was used to tune the ridge penalty (*λ*). Log-spaced candidate values (0.1, 1, 10, 100, 1000, 10000) were evaluated, with predictors standardized using the training set standard deviation. For each participant, region, layer, and unit type, the penalty with the best median performance across folds was chosen. Final performance was quantified as voxel-wise *R^2^*, computed from predictions concatenated across all outer folds.

To separate the contributions of face-, body-, and mixed-selective units, we performed hierarchical variance partitioning. First, we compared face and body units directly by fitting face-only, body-only, and face+body models with the same penalty tuned on face+body. Unique and shared variance components were derived from these fits. Second, we tested whether mixed units added explanatory power beyond face+body by comparing face+body+mixed to face+body, with penalties tuned on the former. The additional contribution of mixed units was defined as the difference between these two models. Penalties of full predictor sets were used to ensure differences in prediction performance reflected differences in predictor sets rather than regularization (Dupré la Tour et al., 2022). Finally, to assess how much variance is shared between face and body selectivity across regions, we calculated the proportion of overlapping variance, defined as the variance jointly explained by face and body units plus the additional variance explained by mixed units, relative to the total variance explained in each region. Because predictors were correlated (especially mixed units with face and body units, respectively), absolute estimates of variance portions should be interpreted cautiously. Nevertheless, the relative magnitudes and ratios across models remain informative, revealing how unit types contribute uniquely and jointly to predicting cortical responses.

To compare encoding performance across unit types while enabling population-level inference, we used a random-effects approach, treating participants as the random factor. For each participant, region, and unit type, we computed the mean *R^2^* across voxels. We then performed paired two-sided *t*-tests on these participant means to compare unit types within each region type (i.e., face-selective, body-selective, and overlap region). To control for multiple comparisons, we applied Bonferroni correction across all pairwise comparisons within each region type.

### Lesioning analyses

To evaluate the functional relevance of different unit types, we used a fixed DNN trained on object recognition. Linear classifiers were trained on the penultimate layer (fc7, post-ReLU) to perform three person perception tasks: face recognition using the VGGFace2 test set (500 identities; Cao et al., 2018), person recognition using the Celeb-ReID training set (632 identities; Huang et al., 2020), and action recognition using the Stanford 40 Actions training set (40 human actions; Yao et al., 2011). For each dataset, images were split into training (60%), validation (20%), and test (20%) sets using stratified sampling over classes. Classifiers were optimized for a maximum of 10 epochs with a batch size of 64. For face recognition, a learning rate of 1×10^-3^ and a weight decay of 1×10^-4^ were used. For person and action recognition, a learning rate of 1×10^-4^ and a weight decay of 1×10^-5^ were used. Task performance was evaluated on the held-out test set with the DNN backbone intact.

To quantify the contribution of different unit types, we lesioned each unit type in the fixed backbone and measured the resulting drop in classifier performance normalized by the baseline performance. For each unit type, the outputs of a matched number of units were set to zero across all layers, and test-set accuracy was recomputed without retraining the classifier. This procedure was repeated for 10 different stratified splits to ensure robustness and to account for potential class imbalances. To compute 95% confidence intervals and perform pairwise comparisons between unit types, we used hierarchical bootstrapping (10,000 iterations), as described below.

### Hierarchical bootstrapping

To account for nested data structures, we used hierarchical bootstrapping to estimate confidence intervals, applying either a two-level or three-level resampling scheme depending on the analysis. For validation analyses aggregating selectivity (*d’*) across models, we used three-level resampling: (i) resampling models with replacement, (ii) resampling layers with replacement within each selected model, and (iii) resampling units with replacement within each selected layer (1,000 iterations). For lesioning analyses aggregating classification performance, we used two-level resampling: (i) resampling cross-validation splits with replacement, and (ii) resampling images with replacement within each selected split (10,000 iterations). In both cases, we computed the statistic of interest across the pooled resample, capturing uncertainty from variability at all levels of the hierarchy.

We used these bootstrap distributions for hypothesis testing, performing one-sample tests to assess whether statistics differed from zero and pairwise tests to compare statistics between conditions. All *p*-values were corrected for multiple comparisons using the Benjamini-Yekutieli FDR procedure (*α* = 0.05), which controls the false discovery rate under arbitrary dependence assumptions (Benjamini & Yekutieli, 2001).

## Data and code availability

The fMRI data supporting these analyses were obtained from the publicly available Natural Scenes Dataset (http://naturalscenesdataset.org/). The Python code (version 3.10.12) used for data analysis and visualization will be made available upon publication of the manuscript.

## Acknowledgments

We thank all members of the VCCN Lab and Daniel Kaiser for their valuable feedback on the manuscript. Special thanks go to Jacob S. Prince for sharing the ImageNet weights for the AlexNet Barlow Twins model and to Stefania Bracci for sharing some of the images used in the validation dataset. L.E.v.D. was supported by a doctoral scholarship awarded by the German Academic Scholarship Foundation. K.D. was supported by the ERC Starting Grant DEEPFUNC (ERC-2023-STG-101117441), the Hessian Ministry of Higher Education, Research, Science and the Arts (LOEWE Start Professorship and Excellence Cluster EXC3066 “The Adaptive Mind”), and the Deutsche Forschungsgemeinschaft (DFG, German Research Foundation, 222641018-SFB/TRR 135 TP). The funding organizations had no role in the study design, data collection and analysis, decision to publish, or preparation of the manuscript.

## Author contributions

L.E.v.D. and K.D. conceived the study. L.E.v.D. carried out the data analysis and wrote the original draft of the manuscript. K.D. reviewed the manuscript and provided critical feedback. K.D. supervised the project.

## Supplementary materials

## Supplementary materials

**Fig. S1.**
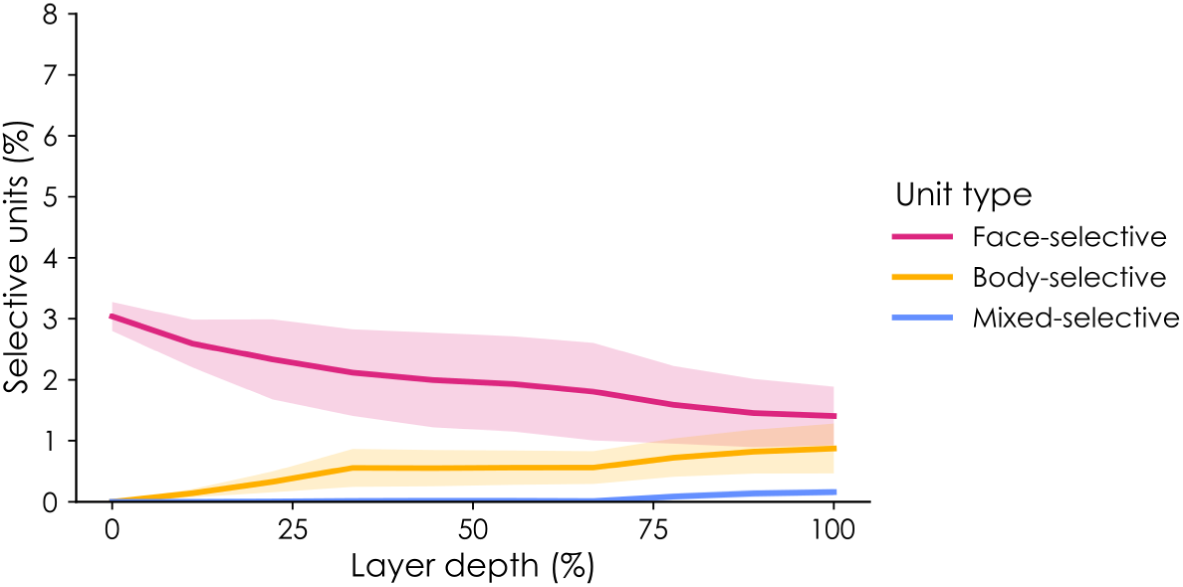
Proportion of selective units in untrained DNNs. Proportion of selective units (%) across normalized layer depth (%), averaged across randomly initialized models (mean ± *SEM*).

**Fig. S2.**
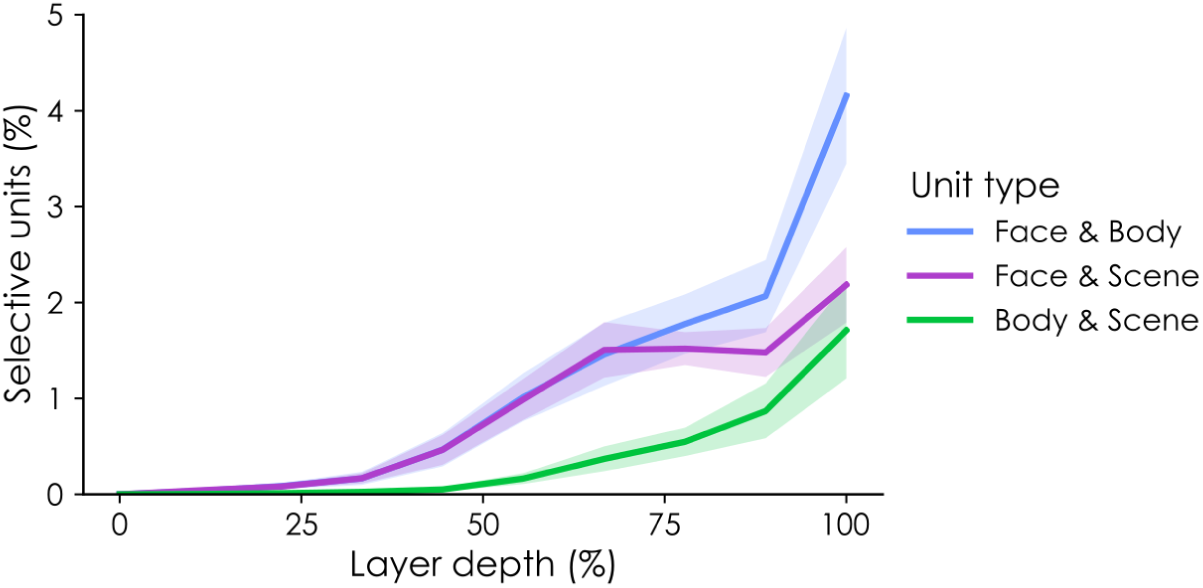
Proportion of mixed-selective units for different category combinations. Proportion of mixed-selective units (%) for different category combinations across normalized layer depth (%), averaged across models (mean ± *SEM*).

**Fig. S3.**
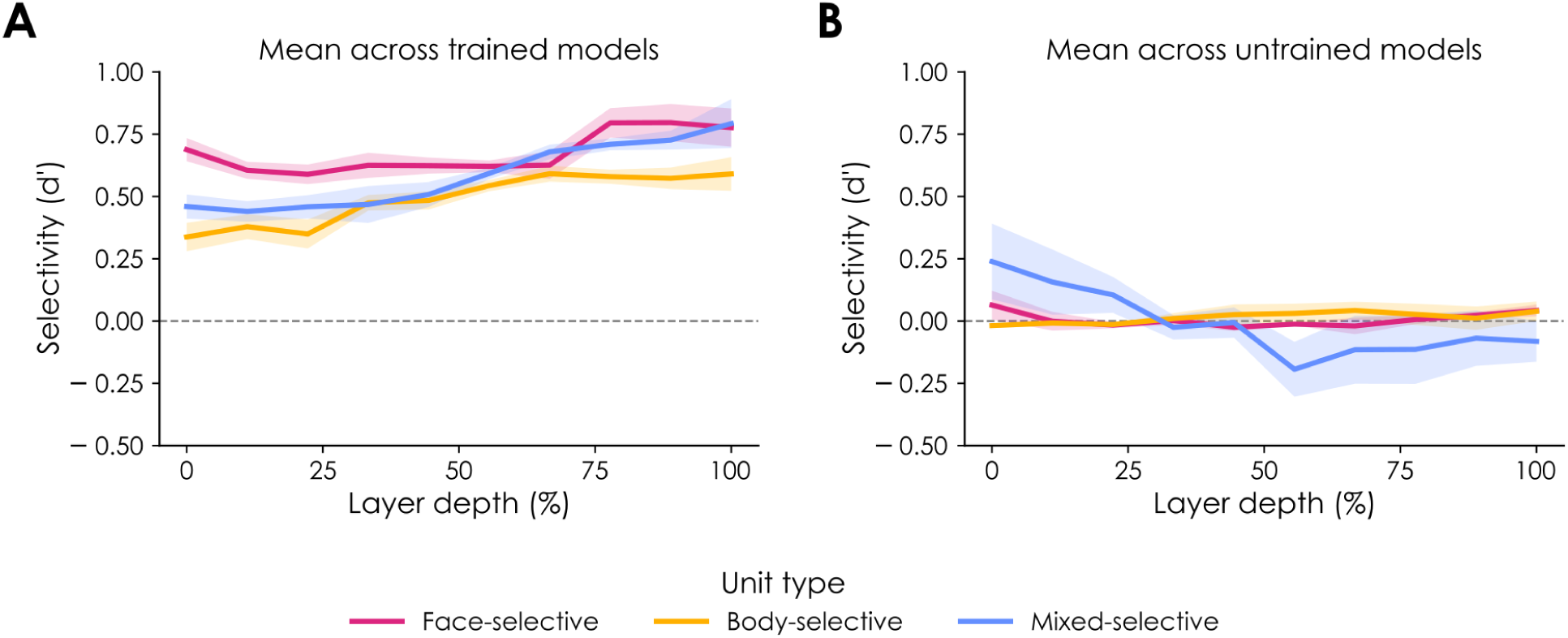
Selectivity across layers. Selectivity (*d’*, preferred category vs. objects) based on the validation dataset for each selective unit type across normalized layer depth (%), averaged across models (mean ± *SEM*). (**A**) Mean across trained models. (**B**) Mean across untrained models.

**Fig. S4.**
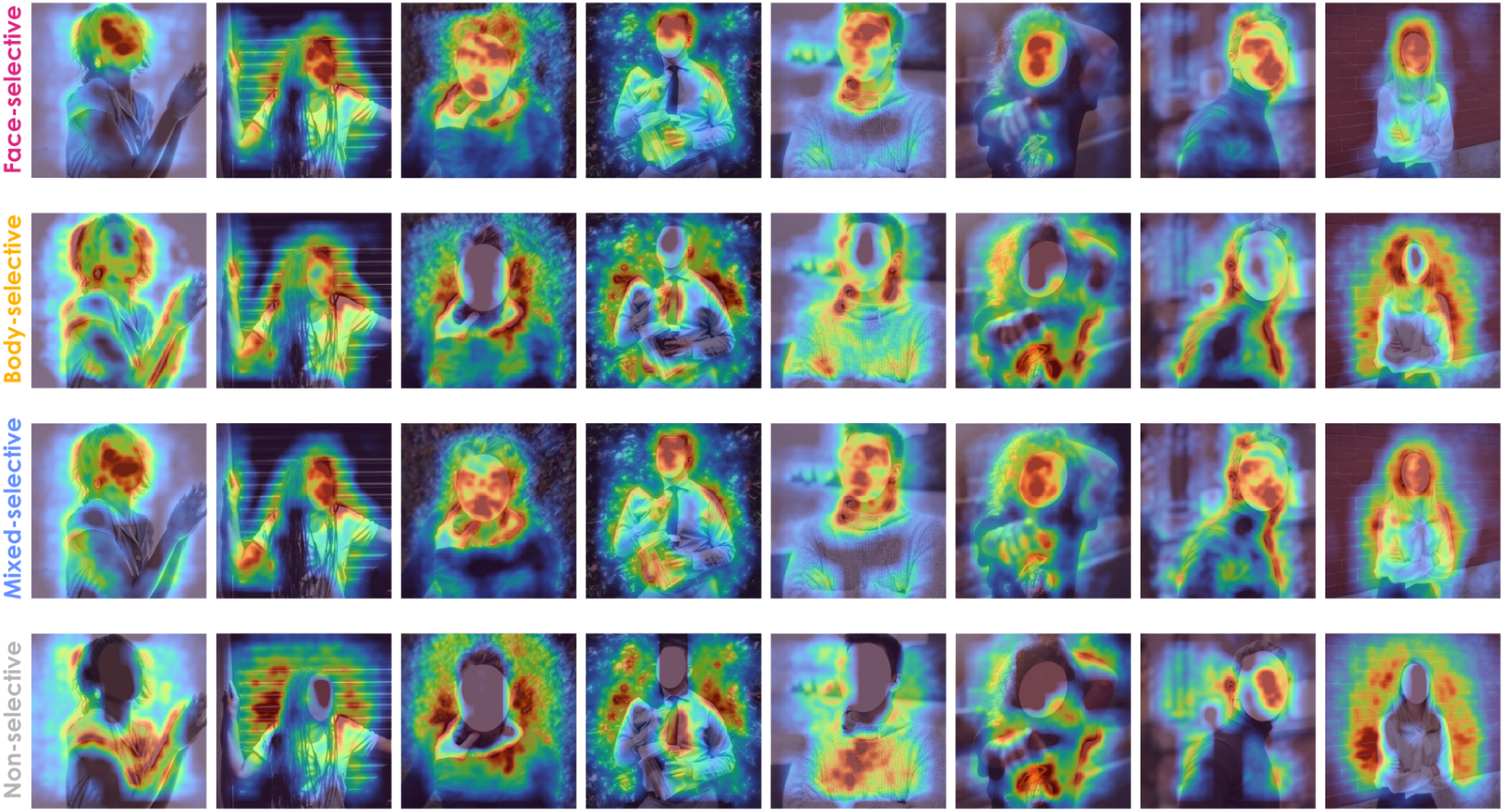
Feature visualizations of each unit type. Guided-GradCAM heatmaps for each unit type in the last convolutional layer of a representative model (supervised AlexNet trained on Ecoset; conv5) overlaid on publicly available example images (source: <u>Pexels.com</u>). Images containing faces were masked afterwards.

**Fig. S5.**
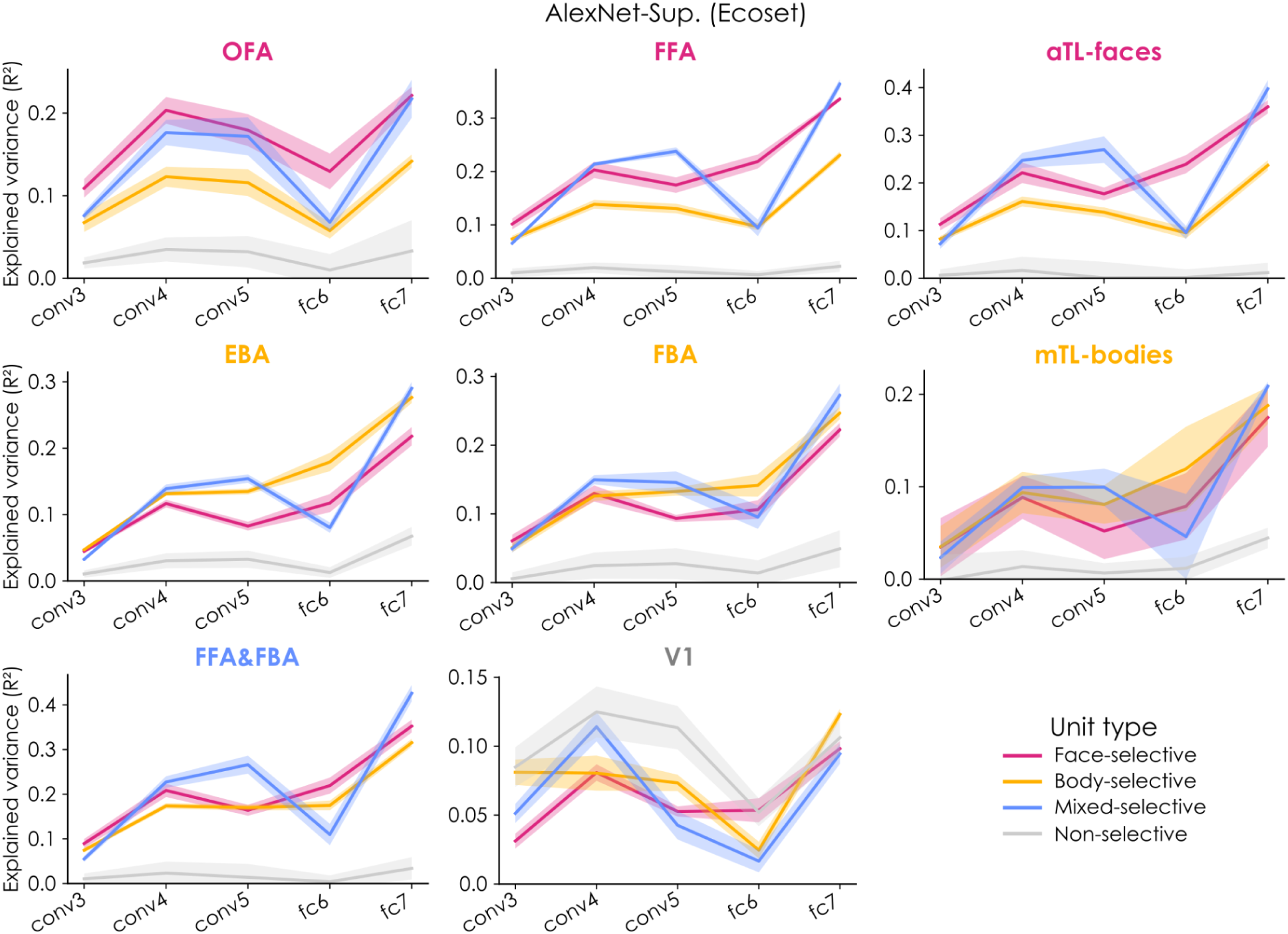
Prediction performance of DNN unit types across layers in different cortical regions. Explained variance (*R^2^*, noise-normalized) of each unit type across layers in each cortical region. Lines and shaded areas show group means and 95% CIs (Cousineau-Morey-corrected).

**Fig. S6.**
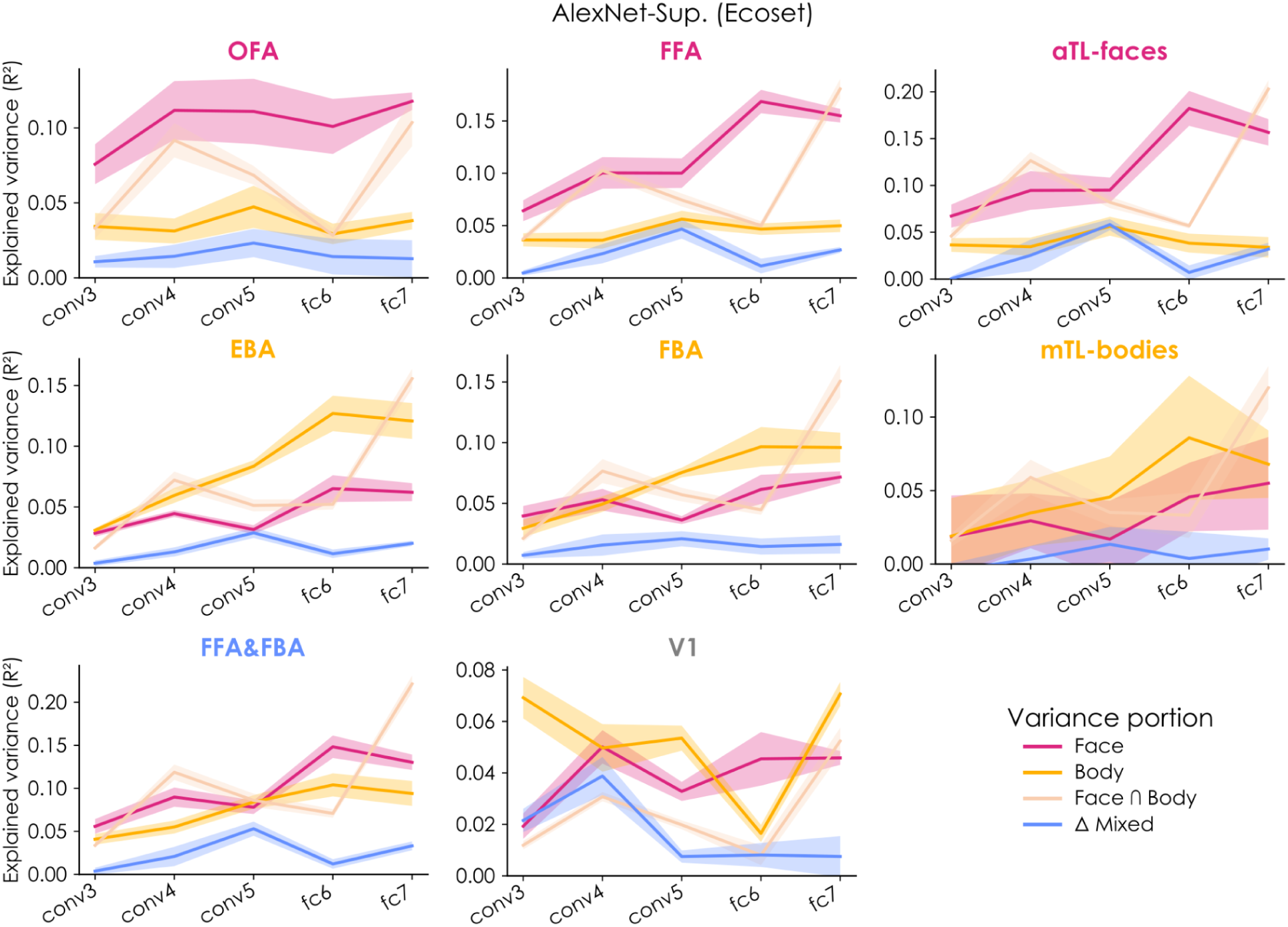
Variance portions of DNN unit types across layers in different cortical regions. Explained variance portions (*R^2^*, noise-normalized) of each selective unit type across layers in each cortical region. Variance partitioning of encoding models fit with multiple unit types showing unique variance explained by face units (Face), unique variance explained by body units (Body), their shared variance (Face∩Body), and the additional unique variance gained by adding mixed units (ΔMixed). Lines and shaded areas show group means and 95% CIs (Cousineau-Morey-corrected).

